# QCT-based computational bone strength assessment updated with MRI-derived ‘hidden’ microporosity

**DOI:** 10.1101/2023.03.30.534902

**Authors:** Samuel McPhee, Lucy E Kershaw, Carola R Daniel, Marta Peña Fernández, Eugenio Cillán-García, Sarah E Taylor, Uwe Wolfram

## Abstract

Microdamage accumulated through sustained periods of cyclic loading or single over-loading events contributes to bone fragility through a reduction in stiffness and strength. Monitoring microdamage *in vivo* remains unattainable by clinical imaging modalities. As such, there are no established computational methods for clinical fracture risk assessment that account for microdamage that exists *in vivo* at any specific timepoint. We propose a method that combines multiple clinical imaging modalities to identify an indicative surrogate, which we term ’hidden porosity’, that incorporates pre-existing bone microdamage *in vivo*. To do so, we use the third metacarpal bone of the equine athlete as an exemplary model for fatigue induced microdamage, which coalesces in the subchondral bone. N=10 metacarpals were scanned by clinical quantitative computed tomography (QCT) and magnetic resonance imaging (MRI). We used a patch-based similarity method to quantify the signal intensity of a fluid sensitive MRI sequence in bone regions where microdamage coalesces. The method generated MRI-derived pseudoCT images which were then used to determine a pre-existing damage (*D*^pex^) variable to quantify the proposed surrogate and which we incorporate into a nonlinear constitutive model for bone tissue. The minimum, median, and maximum detected *D*^pex^ of 0.059, 0.209, and 0.353 reduced material stiffness by 5.9%, 20.9%, and 35.3% as well as yield stress by 5.9%, 20.3%, and 35.3%. Limb-specific voxel-based finite element meshes were equipped with the updated material model. Lateral and medial condyles of each metacarpal were loaded to simulate physiological joint loading during gallop. The degree of detected *D*^pex^ correlated with a relative reduction in both condylar stiffness (*p* = 0.001, R^2^ > 0.74) and strength (*p* < 0.001, R^2^ > 0.80). Our results illustrate the complementary value of looking beyond clinical CT, which neglects the inclusion of micro-damage due to partial volume effects. As we use clinically available imaging techniques, our results may aid research beyond the equine model on fracture risk assessment in human diseases such as osteoarthritis, bone cancer, or osteoporosis.

## 1 Introduction

Sustained periods of cyclic loading below and single overloading events beyond yield stress leads to the accumulation of microdamage in bone (Gupta and Zioupos, 2008; Vashishth, 2007; Wolfram et al., 2011). Initiation and propagation of microdamage serves to dissipate energy to prevent crack coalescence and propagation but causes a reduction in bone stiffness and strength (Gupta and Zioupos, 2008; Seref-Ferlengez et al., 2015), which consequently increases the risk of fracture. In healthy bone, the accumulation of microdamage is regulated by the remodelling process, but if this functional capacity to repair emerging microdamage is suppressed or surpassed, atraumatic stress fractures can occur. Clinically, stress fractures can be divided into fatigue and insufficiency fractures (Datir, 2007). Fatigue fractures arise when the reparative capacity to remove accumulated microdamage is surpassed and are most commonly encountered in athletes and military personnel where demanding training regimes are the primary predisposing factors to bone fracture (Wentz et al., 2011). Insufficiency fractures, on the other hand, occur during physiologic loading of mechanically compromised bone that is weakened either by low mass, quality, or impaired reparative function. In instances where weakening is driven by a reduction in mass, for example in osteoporosis or osteolytic bone metastasis, the weakening is further compounded by the accumulation of microdamage (Atkins et al., 2019). Ageing is a major risk factor for diseases that increase bone fragility, which, when combined with the inherent increase in microdamage accumulation in aged individuals (Burr et al., 1997), makes monitoring of microdamage *in vivo* an important clinical objective. Besides the impractical means of obtaining biopsies for histological investigations, microdamage detection *in vivo* remains an unsolved clinical challenge (Chapurlat and Delmas, 2009) due to the micro-and nanoscale size order. *In vivo* monitoring of accumulating bone microdamage would improve current fracture risk diagnostics and potentially allow for preventative interventions prior to fracture especially in such cases that are not accompanied by a loss of bone mass, for example, atypical femoral fractures (Ettinger et al., 2013; Starr et al., 2018).

Classically, fracture risk assessment in a clinical setting is done by measuring areal bone mineral density by dual-energy X-ray absorptiometry (Bates et al., 2002). While areal bone mineral density is a good predictor of fracture risk, it is a singular measure of bone quality which does not allow assessment of structural determinants of bone strength. The short-comings of dual-energy X-ray absorptiometry are evidenced by the failure to capture over half of non-vertebral fractures that occur in aged individuals with normal bone mineral density (Schuit et al., 2004). Further improvements have been made to clinical fracture risk assessment by incorporating geometric and mineral density distribution information from quantitative computed tomography (QCT) scans into finite element models (Adams et al., 2018; Chevalier et al., 2008; Dall’Ara et al., 2012; Keaveny et al., 2020; Luisier et al., 2014). The achievable voxel size of clinical QCT is still greater than the typical length scale of microcracks in bone (Arlot et al., 2008a), and as such, QCT-derived finite element modelling falls short of incorporating accumulated microdamage.

Where these X-ray attenuation-based imaging modalities visualise the mineralised component of bone, magnetic resonance imaging (MRI), visualises the adjacent soft tissues and bone marrow. MRI is sensitive to pathophysiological changes in marrow composition, which present as alterations in signal intensity. This sensitivity lends the technique to early detection of chronic and acute traumatic bone injuries which may be otherwise undetectable by plain radiographs or CT (Datir, 2007). In the context of traumatic bone injury, the primary imaging characteristic is an increased signal intensity on fluid sensitive T2w and STIR se-quences (Linn et al., 2009; Molfetta et al., 2022), which reflects the presence of underlying edema, haemorrhage, hyperemia, and trabecular microdamage as evidenced by histological studies (Matheny et al., 2017a; Muratovic et al., 2018; Rangger et al., 1998; Taljanovic et al., 2008). Qualitative MRI-based grading of bone stress injuries exists (Fredericson et al., 1995; Kiuru et al., 2003), however, if the signal alterations could be reliably quantified, it could be used to supplement existing density-based fracture risk modelling techniques with quantitative information on bone quality.

MRI signal hyperintensity in bone is a nonspecific imaging characteristic that is attributable to multiple bone pathologies (Maraghelli et al., 2021). For this reason, we require a suitable model where the mechanism of injury is consistent and understood. We here utilise the equine athlete as an exemplary model for stress fractures. Sustained periods of intense high-speed exercise during training and racing induces subchondral microdamage in bones that comprise the lower limbs (Muir et al., 2008, 2006; Shaffer et al., 2022a; Whitton et al., 2018). The most common and fatal fracture events in Thoroughbred racehorses involve the metacarpophalangeal joint (Parkin et al., 2004). Here, the distal end of the third metacarpal bone is supported by the proximal phalanx and proximal sesamoid bones at oblique angles to the long axis of the metacarpal bone. The phalanx-metacarpal joint reaction force is well tolerated due to the large area and high congruency between the articular surfaces (Harrison et al., 2014; Riggs et al., 1999). The impaction of the proximal sesamoid bones with the palmar aspect of the condyles of the distal metacarpal bone, however, leads to highly consistent morphological changes and fracture configurations (Stover, 2017). The repetitive high-strain environment in the subchondral bone suppresses resorption (Holmes et al., 2014), driving rapid apposition of matrix material, resulting in marrow space infilling and loss of structural anisotropy (Boyde, 2003; Boyde and Firth, 2005; Rubio-Martínez et al., 2010). This additive response increases subchondral bone volume fraction to levels comparable with cortical bone (Malekipour et al., 2018; Martig et al., 2018; Riggs et al., 1999; Rubio-Martínez et al., 2008; Whitton et al., 2010). While increasing bone volume fraction usually increases bone strength and fatigue resistance (Martig et al., 2020; Rapillard et al., 2006), there is an association between densification of the subchondral bone in the palmarodistal aspect of the metacarpal condyles and condylar fracture (Peloso et al., 2019; Tranquille et al., 2017; Whitton et al., 2010). This modelling dominant response, when coupled with further athleticism, leads to the accumulation of microcracks in the densified subchondral bone of horses in training (Muir et al., 2008; Whitton et al., 2018). The consequence of this accumulated microdamage is an uncharacteristically low apparent stiffness despite exhibiting a high bone volume fraction. For example, Martig et al. (2020) reported for subchondral bone volume fraction over 0.85, stiffness values around 2.3 GPa. At this amount of bone volume fraction, one would expect stiffness values of 18 GPa (Mirzaali et al., 2016; Wolfram and Schwiedrzik, 2016) and it could be inferred that bone strength is similarly affected. It is known that bone volume fraction is the main determinant of both stiffness and strength (Currey, 2002; Maquer et al., 2015; Musy et al., 2017) and this associative loss of bone quality, despite drastic adaptation, affirms that exercise-induced microdamage accumulation has a critical impact on the load-bearing capacity and consequently fracture incidence, which cannot be detected by CT-based diagnostic means.

MRI is routinely used as a diagnostic tool for elucidating musculoskeletal injury in Thoroughbred racehorses. If an imaging characteristic that is indicative of microdamage can be identified and therein quantified in terms of metacarpal bone mechanical competence, these diagnostic imaging protocols, including image-based computational models, could be used to inform clinicians as to the associated risk of continued exercise activity. From a translational perspective, such imaging characteristics could be helpful for the diagnosis of existing microdamage in humans *in vivo*. Therefore, our aims were to (i) identify a suitable imaging modality that may be characteristically indicative of a damage state in subchondral bone; (ii) identify a suitable method to quantify MRI signal hyperintensity; (iii) use this quantification to derive a pre-existing damage variable attributed to a damage state within the distal metacarpal subchondral bone; (iv) establish a QCT-derived bone volume fraction based finite element modelling protocol for *in silico* strength tests of the distal metacarpal condyles and; (v) update these image-based finite element models with the MRI-derived pre-existing damage variable to evaluate the relative reduction in whole bone stiffness and strength.

## 2 Materials and Methods

### 2.1 Samples

Ten forelimbs (5 left, 5 right) from five Thoroughbred racehorses (geldings, aged 3-11 years) were included in the study. Each limb was disarticulated at the antebrachiocarpal joint within 24 hours post-mortem and frozen at −20 °C until required for imaging. Before an imaging session, the limbs were rested at room temperature for 24 hours to ensure that they were completely thawed. We then imaged the limbs immediately thereafter with an initial radiograph to screen for metallic objects.

### 2.2 Imaging

The distal forelimbs of the horses were scanned by clinical CT (SOMATOM Definition AS, Siemens Healthineers, Germany; voltage: 120 kV, current: 80 mA, matrix size: 512 x 512, slice thickness: 0.6 mm) and reconstructed with a voxel size of 0.273 mm x 0.273 mm x 0.4 mm. Each limb was scanned with a hydroxyapatite phantom (model number 8783219, Siemens, Germany) to accommodate calibration to a bone mineral density scale (QCT).

We then scanned each metacarpophalangeal joint at 3 T (Siemens MAGNETOM Skyra, Siemens Healthineers, Germany) using a 15-channel knee coil. The imaging protocol included a T1-weighted TSE (Turbo spin-echo) sequence acquired in three orthogonal planes (repetition time (TR) = 1660 ms, echo time (TE) = 20 ms, 3 averages, voxel size = 0.3 mm x 0.3mm x 1.6 mm) and an STIR sequence acquired in the sagittal and coronal planes (TR/TE = 9930/44 ms, inversion time (TI) = 220 ms, voxel size = 0.5 mm x 0.5 mm x 1.5 mm).

### 2.3 QCT Image processing

The following QCT image processing was conducted in ImageJ (NIH ImageJ 1.53 (Abràmoff et al., 2004)). We parsed the raw 16-bit grey-level QCT images to a bone mineral density (BMD) scale by linear transformation based on the hydroxyapatite phantom. We then cropped and resampled the BMD images to isotropic voxels with a size of 0.3 mm by B-Spline interpolation. Binary masks of the metacarpal bones were then generated. To do so, we used 3D median filtering (kernel size = 2) to remove noise before applying a contrast-based local thresholding filter (kernel size = 50) to convert the image to binary. We then removed background voxels using a connectivity filter, followed by a flood fill filter to produce a continuous solid volume, representative of each metacarpal.

#### 2.3.1 BMD to BVTV calibration

The spatial resolution of QCT is in the size order of single trabeculae plus marrow space, preventing identification of bone-marrow-interfaces. Therefore, it is not possible to quantify bone volume fraction (BV/TV) by segmentation techniques. Dall’Ara et al. (2013) demon-strated, however, that QCT-derived BMD is linearly correlated to µCT-derived BV/TV. We used a calibration law presented by Dall’Ara et al. (2013) to transform BMD to BV/TV:

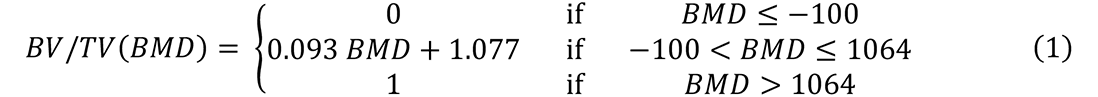

The calibration law is derived from *ex vivo* µCT and clinical QCT scans of human femurs. Nevertheless, we deem it applicable to the equine metacarpal based on µCT evidence pre-sented by Martig et al. (2018), who extracted subchondral bone cylinders from horses in training and showed that with volume fractions comparable to cortical bone (>0.95), the corresponding tissue mineral density is in the range of 0.92-1.08 gHA/cm^3^ (Martig et al., 2018).

### 2.4 MR image processing

#### 2.4.1 Dense bone volume segmentation

We used an automated super-resolution reconstruction algorithm (NiftyMIC (Ebner et al., 2020)) to reconstruct isotropic images (voxel size = 0.3 mm) of the T1w and STIR sequences from their respective orthogonal image stacks. We then registered the isotropic T1w and STIR images to the QCT images. For each limb, we first manually aligned the T1w image to its respective QCT image using a linear transformation in Slicer 3D (Slicer 4.11 (Fedorov et al., 2012)). We then registered the T1w image to the QCT image by affine registration, using a mutual information metric (histogram bins = 32) (SimpleElastix (Marstal et al., 2016)). The resultant affine transform matrix was used to transform and register the STIR image to its respective QCT image.

The volume and depth of dense subchondral bone are associated with increased incidences of condylar fracture (Tranquille et al., 2017; Whitton et al., 2010). We calculated the volume of this dense subchondral bone to investigate its influence on whole bone strength of each metacarpal bone. The dense subchondral bone was segmented by one investigator (SM) and verified by a trained clinical investigator who regularly examines clinical MRI in the context of equine musculoskeletal injury (SET). The segmentation criterion was chosen as any continuous T1w hypointense volume subjacent to the articular surface in the palmar aspect of the lateral and medial condyles (Fig. 1d). We calculated a dense bone volume fraction (DBV/MV) (Peloso et al., 2019) for each bone by dividing the segmented dense bone volume (denoted as DBV) by an anatomically consistent volume of the distal end of the metacarpal (denoted as MV). Following Peloso et al. (2019), we chose the anatomical landmark as a transverse plane that intersected the proximal palmar aspect of the distal metacarpal, where the curvature of the subchondral bone plate changes from a convex to a concave shape (Fig. 1). The distal metacarpal was then segmented into a lateral and medial side by a sagittal plane through the sagittal ridge.

**Fig. 1.**
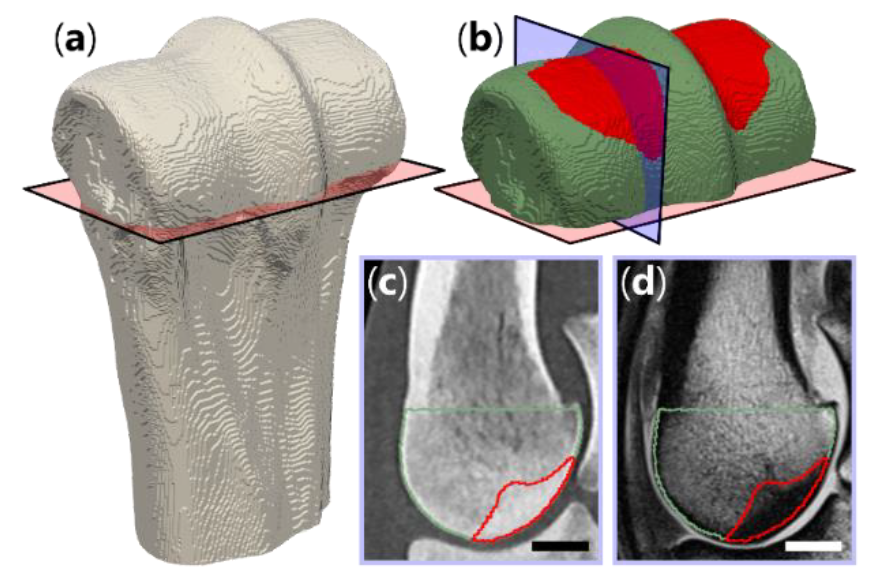
Dense bone segmentation: (a) Distal end of the third metacarpal bone where the red transverse plane is used to normalise the volume of subchondral dense bone using an anatomically normalised volume of the distal metacarpal (MV). (b) The red is the segmented dense bone (DBV), and the green is the healthy bone that comprises MV. The blue sagittal plane here is an arbitrary slice corresponding to a QCT (c) and T1w (d) sagittal slice of the distal third metacarpal where the red and green lines bound the dense bone and healthy bone, respectively. The scale bar here is 10 mm.

#### 2.4.2 Intensity-based pseudoCT generation

To quantify relative signal differences in the dense bone, we employed a patch-based (cuboidal subvolumes) similarity search method to generate MRI-derived pseudoCT (pCT) images. Andreasen et al. (2015) present a patch-based pCT generation methodology and we herein outline their methodology. The methodology relies on a registered CT-MR dataset, from which a test MR image can be queried.

First, the dense bone segmentation (Section 2.4.1) of the T1w image (Fig. 2a-i) was used to divide the QCT (Fig. 2a-ii) and STIR (Fig. 2a-iii) images into healthy and dense bone sub-regions (Fig. 2b and c). This allowed for intra-sample comparison which was chosen to avoid the need for image standardisation that is required when considering quantitative inter-sample techniques (Nyul et al., 2000).

**Fig. 2.**
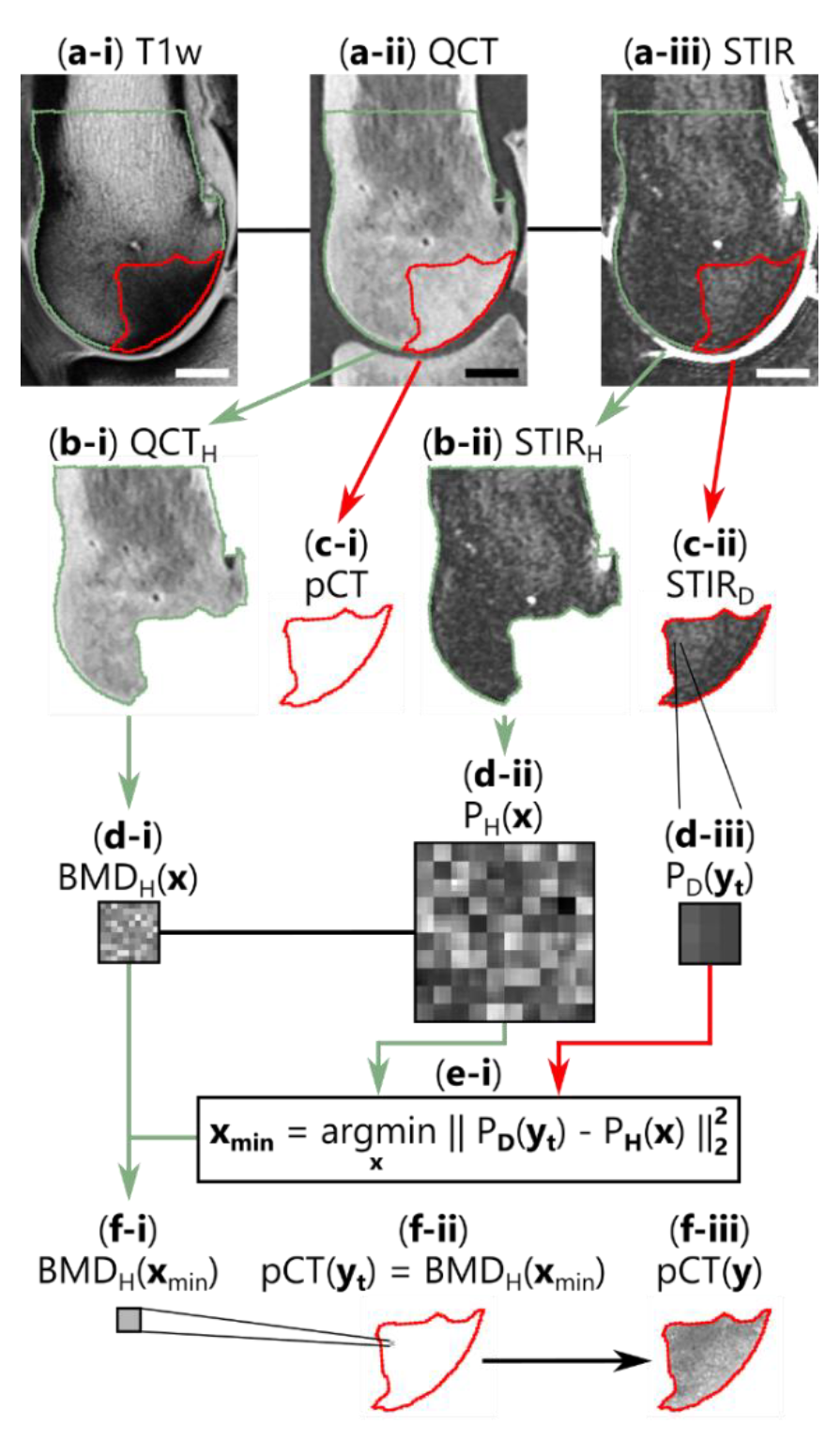
PseudoCT generation pipeline: (a) Registered T1w (a-i), QCT (a-ii) and STIR (a-iii) images. Scale bar is 10 mm. (b-c) The QCT and STIR images are segmented into a healthy subregion (subscript H, b-i and b-ii) and dense bone subregion (subscript D, c-i and c-ii). (d-e) A patch database is generated from STIRH subre-gion (d-ii, 10 x 10 array of 2D 3 x 3 patches for illustration) along with a spatially aligned BMD database derived from the QCTH subregion (d-i, 10 x 10 array of BMD values for illustration). A test patch (d-iii) from the STIRD subregion is queried against database STIRH. The spatial position of the voxel in STIRH that mini-mizes the squared Euclidean distance (e-i) to the test patch is used to assign a BMD value (f-i) to the pCT (f-ii). All voxels that comprise STIRD are queried until the pCT is ‘filled’ (f-iii).

We generated a database of patches, *P_H_*(***x***), for all voxels that comprise the healthy sub-region in the STIR image (Fig. 2d-ii), where a patch consisted of intensity values extracted from a 3 × 3 × 3 voxel subvolume centred about its respective spatial position, ***x***. As the QCT and STIR images were registered, any STIR-derived patch with spatial position ***x*** in database, *P_H_*(***x***), had a corresponding BMD value, *BMD_H_*(***x***), from the aligned QCT image (Fig. 2d-i).

A patch database for the dense bone subregion, *P_D_*(***y***), was similarly generated from the STIR image. Similarity between each patch in *P_D_*(***y***) was iteratively compared against the healthy patch database *P_H_*(***x***). During one iteration, a single test patch with position ***y_t_*** was queried (Fig. 2d-iii). We chose the squared Euclidean distance, *d*(***x***, ***y***), (Fig. 2e-i) as a simi-larity metric:

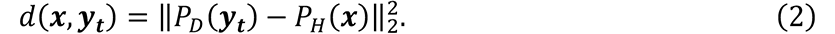

A nearest neighbour search using a kd-tree algorithm (sklearn.neighbors.KDTree (Pedregosa et al., 2011)) was used to return the spatial position, ***x***_min_, of the patch in *P_H_*(***x***) that minimised *d*(***x***, ***y***) where:

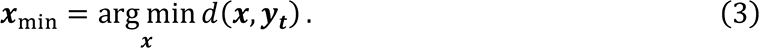

Prior to any voxels being queried, a zero filled pCT image with consistent dimensions as the dense bone subregion was generated as, *pCT*(***y***) (Fig. 2f-ii). Then for each iteration, a BMD value was assigned to the pCT image (Fig. 2f-i and f-ii) by:

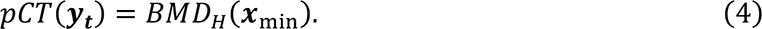

Each voxel that comprised the dense bone subregion, *P_D_*(***y***), was queried and the corresponding bone mineral density was assigned until the pCT image, *pCT*(***y***), was ’filled’ (Fig. 2f-iii). The segmented dense bone subregions of all ten metacarpals were queried.

We then excluded the dense bone subregion and repeated the procedure where the lateral side of the distal metacarpal was queried against the medial side. We then computed a histogram intersection match score between the QCT histograms, *H_QCT_*, and pCT histograms, *H*_*pCT*_, by (Swain and Ballard, 1991):

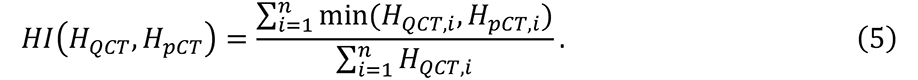

Here, *n* is the number of histogram bins and *HI*(*H_QCT_*, *H*_*pCT*_) is a fractional match value between 0 and 1. We consider an intersection match score of 0.9 sufficiently similar. The pCT similarity search algorithm was implemented in python (3.6). Each query was parallelised using a python binding for the message passing interface (mpi4py (Dalcín et al., 2005)), utilising 32 CPUs on Heriot-Watt University’s HPC facility ‘Rocks’.

#### 2.4.3 MRI-derived damage surrogate

To quantify the underlying microdamage in the subchondral bone of the palmar aspect of the distal metacarpal bone, we introduce a pre-existing damage parameter, *D*^pex^, derived from the quantification of STIR signal intensity in Section 2.4.2. The signal hyperintensity in the fluid sensitive STIR sequence is a consequence of a wide range of reparative events occurring in the subchondral bone unit. For this reason, we refrain from interpreting our quantification explicitly as damage. Instead, we refer to it as a damage surrogate, whereby the signal increase represents an underlying damage state or state of porosities such as micro-damage and increased vasculature which is unobservable by clinical CT methods. Mechanistically, this state manifests as a volume density of micron and submicron cracks and cavities which we denote *D*^pex^.

We derive *D*^pex^ by following Lemaitre (1984), where we consider a representative volume element *V* that can be imaged by both CT and MRI. We consider *V* to be large enough to contain many defects but small enough to be a material point. This assumption is justified by the resolution of the used imaging modalities that achieve a voxel size of 300 µm, whereas microcracks accumulated through repetitive loading are in the size order of 10 µm (Arlot et al., 2008a; Larrue et al., 2011; Wolfram et al., 2016). Lemaitre (1984) then defined damage based on a surface density of microcavities on a plane that transects *V* and its surface normal to define a geometric damage variable. For simplicity, we assume isotropy such that the *D*^pex^ is defined by a loss of effective carrying volume by:

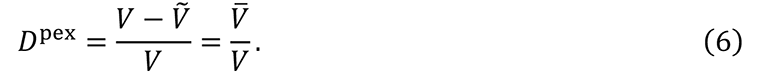

Here, 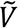 is the effective solid phase volume and 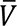 is the volume of microcavities that occupies total volume *V*. To determine *D*^pex^, we utilise the quantification of STIR signal intensity in segmented dense bone region (Section 2.4.2), where we know that microdamage accumulates (Muir et al., 2006; Whitton et al., 2018). The QCT imaging mode provides a means to determine BV/TV and the STIR-derived pCT provides a fluid dependant apparent bone volume fraction (BV/TV_pCT_). Since the QCT and pCT are registered, both images can be normalised to the same representative volume element. By utilising a rule of mixture, we then approximate that a loss of apparent bone volume fraction in the pCT is attributed to fluid filled microcracks and cavities where:

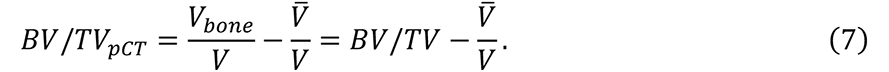

Substituting Eq. (7) into Eq. (6) allows us to redefine our damage variable *D*^pex^ as:

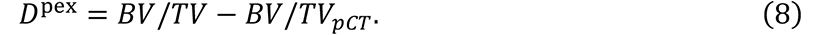

We calculate BV/TV and BV/TV_pCT_ for the segmented dense bone region in lateral and medial side of each metacarpal by calibrating its respective median BMD value (Fig. 4e-f) to BV/TV using Eq. (1).

### 2.5 Finite element modelling

#### 2.5.1 Material model

We adopted the constitutive model proposed by Schwiedrzik and Zysset (2013) in this study. To capture the post-yield behaviour of bone, we utilise a quadric yield surface (Schwiedrzik et al., 2013) and we refer to Schwiedrzik and Zysset (2013) for the numerical implementation of the stress return algorithm and herein briefly outline the changed constitutive equations. The following notations are used, scalars are denoted as standard minuscules or majuscules (*x* or *X*), vectors are denoted as bold-face minuscule (***x***), second order tensors denoted as bold-face font majuscules (***X***), and forth order tensors are denoted as scripted majuscules (𝕏). The ⊕ symbol is the tensor product and 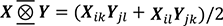 is the symmetric product of two second order tensors. ***X***_*ij*_ = 𝕐_*ijkl*_***Z***_*kl*_ is the trans-formation of a second-order tensor with a fourth-order tensor and ′:′ denotes the double contraction of two tensors.

Small deformations are assumed such that total strain, ***E***, can be split additively into elastic and plastic strains (Green and Naghdi, 1965):

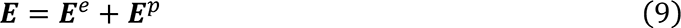

The free energy potential for the material model is then formulated of an isotropic stiffness tensor, 𝕊, and a scalar damage variable, *D*, by:

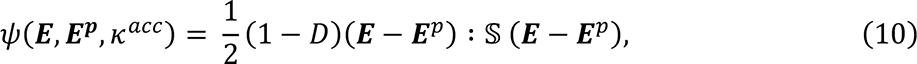

then the stress, ***S***, is derived by:

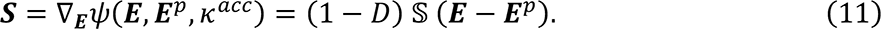

The stiffness tensor, 𝕊, was chosen as isotropic and takes the form (Gross et al., 2013):

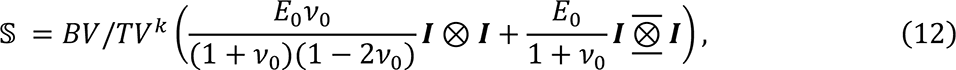

where, *E*_0_ and *v*_0_ are the elastic modulus and Poisson’s ratio of a pore-less material with at least cubic symmetry. *BV/TV^k^* is a power density tissue function that scales all components of a respective tensor by incorporating BV/TV. The influence of the tissue function on stiffness and compressive and tensile strength is shown in Fig. S1.

The scalar damage variable, *D*, is a state internal variable that represents a volume density of micron-and submicron voids which reduces all components of the stiffness tensor. It is derived from plastic strain history and is defined as a function of accumulated plastic strain, *κ^acc^*, by (Charlebois et al., 2010):

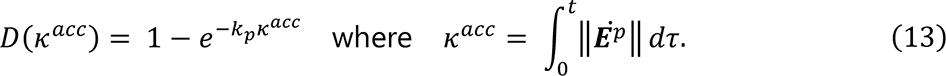

Mirzaali et al. (2015) investigated loading mode dependent damage accumulation in osteonal bone and observed that inducing microdamage through compressive overloading reduced both the stiffness and strength when reloading in tension. Mirzaali et al. (2015) proposed a damage formulation that incorporates this existing overloading history. We borrow this idea of an existing deterioration to capture the mechanical degradation imposed by the increased density of subchondral microcracks in the metacarpal condyles which is induced by high strain exercise (Whitton et al., 2018). While microcrack families are loading mode dependent (Wolfram et al., 2016), for simplicity, we incorporate this overloading history as an isotropic impact by updating the damage formulation as:

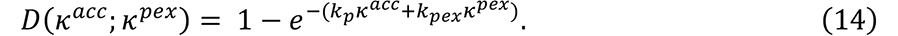

Here, *κ^pex^* implies that there is pre-existing accumulated plastic strain responsible for the nucleation and accumulation of a pre-existing subchondral microcrack density. We postulate that these resemble the aforementioned damage state such that we can derive κ^pex^ from our MRI-derived damage surrogate, *D*^pex^ (Section 2.4.3) as:

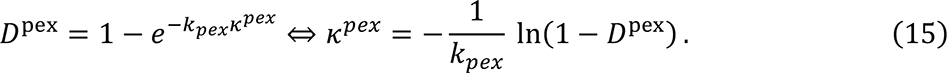

The coefficient *k_pex_* allows us then to weigh the actual influence, however as we currently lack suitable experimental data, we assume this to be *k_pex_* = 1.

The pre-existing accumulated plastic strain then affects the yield criterion. We chose a quadric criterion proposed by Schwiedrzik et al. (2013):

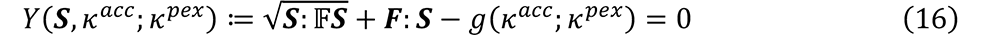

The fourth-order and second-order tensors, 𝔽 and ***F***, control the origin, shape, and orientation of the yield surface. Following Schwiedrzik et al. (2013) these tensors are:

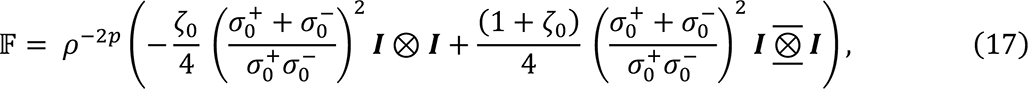

and:

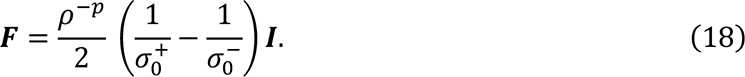

Here, 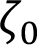, 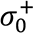, and 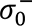 are an interaction parameter which governs the shape of the surface and the uniaxial yield stress in tension and compression respectively. *g*(κ^acc^; κ^pex^) in Eq. (16) is an isotropic exponential hardening function:

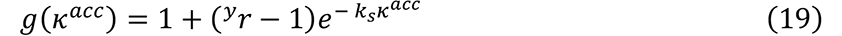

Here, *k_s_* is an experimentally derived exponent and ^*y*^*r*, is the yield-to-strength ratio. Mirzaali et al. (2015) present an exponential hardening function with an exponential decay to account for a reduction in tensile strength in the presence of compressive plastic strain history. While they show conversely that tensile overloading has negligible impact on compressive strength, for simplicity we introduce an isotropic contraction of the yield surface without affecting the post-yield behaviour by:

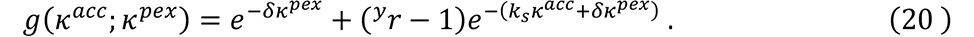

Herein, δ is a parameter that controls the effect of the pre-existing weakening on the initial strength. Mirzaali et al. (2015) introduced this parameter to capture the impact of a compressive overload on a subsequent tensile loading. Our data, however, did not allow identifying this parameter as specialised experiments similar to those by Mirzaali et al. (2015) were not possible. We assume this parameter to be *δ* = 1.

#### 2.5.2 Model parameterisation

We parameterised the model based on Les et al. (1994), who conducted uniaxial compression of 350 bone cylinders extracted from equine third metacarpal bones. Prior to testing, each specimen was scanned and power relationships between mechanical properties and BMD are presented. We use the calibration in Eq. (1) to convert their presented BMD values to BV/TV. Direct interpretation of the power law regression fitting parameters results in an ultimate compressive strength, 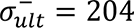 MPa, for a bone volume fraction, BV/TV = 1. This compressive strength is comparable to other studies conducted on equine (Riggs et al., 1993; Tüfekci et al., 2014) as well as osteonal bovine and human bone (McCalden et al., 1993; Mirzaali et al., 2016; Wolfram and Schwiedrzik, 2016). Similarly, the regression fitting parameters resulted in an elastic modulus, *E*_0_ = 15.1 GPa, for a bone volume fraction, BV/TV = 1, which is comparable to some studies conducted on equine bone (Bigley et al., 2008; Les et al., 1997; Riggs et al., 1993). However, this modulus is relatively low when compared to other studies on equine (Gibson et al., 2006) and human bone (Bayraktar et al., 2004; Mirzaali et al., 2016, 2015). This may be explained by the authors use of a platens test, where end-artifact errors produce systematic underestimations in modulus measurements (Keaveny et al., 1997). Nevertheless, we used the modulus presented by Les et al. (1994) as it was the best available data that captured the full physiological range of bone mineral density. We chose an ultimate tensile strength of 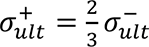 to account for asymmetry in tensile and compressive strength based on observations by Wolfram et al. (2012) and Mirzaali et al. (2015,2016). Finally, the Poisson’s ratio, *ν*_0_, and interaction parameter, 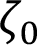, were chosen from Gross et al. (2013) and Panyasantisuk et al. (2016) respectively. The parameters are summarised in Table 1.

**Table 1.** – Elasticity and strength parameters used in the constitutive law that captures the mechanical behaviour of bone tissue (Schwiedrzik et al., 2013; Schwiedrzik and Zysset, 2013). Parameters taken from Les et al. (1994), Gross et al. (2013) and Panyasantisuk et al. (2016) are denoted by (*), (+) and (-) respectively.

| Elasticity |  |  | Strength |  |  |  |
| --- | --- | --- | --- | --- | --- | --- |
| $E_0$ | $\nu_0$ | $k$ | $\sigma_{ult}^+$ | $\sigma_{ult}^-$ | $\zeta_0$ | $p$ |
| in MPa | in - | in - | in MPa | in MPa | in - | in - |
| 15100 (*) | 0.246 (+) | 2.25 (*) | 136 (*) | 204 (-) | 0.280 (+) | 1.79 (*) |

#### 2.5.3 Finite element modelling

The binary mask of each metacarpal was down sampled by a factor of two and converted directly to linear hexahedral elements. For any given element, the mean BMD was computed for a cubic volume of interest (edge length = 1.5 mm) centred about its element centroid (Dall’Ara et al., 2011; Luisier et al., 2014). The element specific BMD was then converted to BV/TV (Fig. 3b) using calibration Eq. (1). Peña Fernández et al. (2022) show that binning a continuous distribution of mineral densities sufficiently captures bone tissue heterogeneity. We chose to similarly bin the elements into sets depending on bone volume fraction (bin size = 0.05; 20 bins). The segmentation of the dense bone region (Section 2.4.2) was used to assign the limb specific magnitude of *D*^pex^ to elements that comprised this region.

**Fig. 3.**
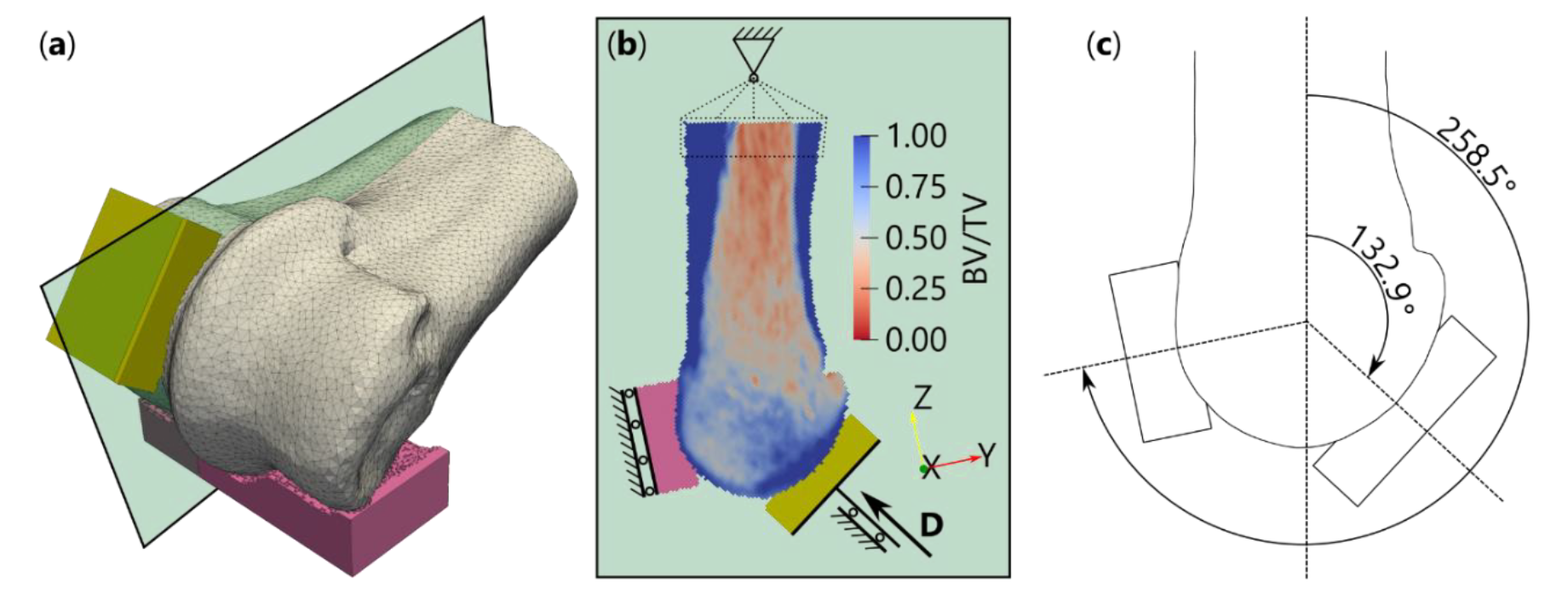
Finite element model setup: (a) Illustration of the congruent mesh blocks used to constrain and compress the distal metacarpal condyles. The orange block on the palmar aspect of the condyle is repre-sentative of a proximal sesamoid bone, whereas the pink block represents the proximal phalanx. The mesh has been smoothed for illustrative purposes. The green sagittal plane corresponds to the slice shown in (b); which is an example of the BV/TV distribution of the distal metacarpal. The boundary conditions are illus-trated as follows; the proximal end of the metacarpal was tied to a node with translational degrees of free-dom constrained; the proximal phalanx mesh block translation was constrained in the y-direction and rota-tion was constrained about the z-axis; the translation of the sesamoid bone was constrained in the x-direc-tion and a displacement, D, was applied and an angle depicted in (c), showing the metacarpal-sesamoid extension angle of 132.9° and the metacarpal-phalanx extension angle of 258.5°.

We chose boundary conditions to reflect a hypothetical *ex vivo* testing assembly informed by joint alignment data (Dall’Ara et al., 2013; Grassi et al., 2020; Kok et al., 2021; Varga et al., 2016). Congruent mesh blocks were added to the palmar and dorsal aspects of the distal metacarpal to simulate contact with the proximal sesamoid bones and proximal phalanx, respectively (Fig. 3a). The chosen angles at which the blocks intersected the condyles were interpreted from data presented by Shaffer et al. (2022b), who measured the motion of the metacarpophalangeal joint under varying physiologically representative axial loads applied the metacarpal. They show that under gallop, the metacarpal-phalanx extension angle is approximately 258.5° (Fig. 3c). They present a relative extension angle change of the proximal sesamoid bones from stand to gallop of approximately 26°. The positioning of the excised limbs during image acquisition in this study prevented direct assessment of the joint alignment during stand. To approximate the relative position of the proximal sesamoid bones with respect to the third metacarpal bone during stand, we used low field standing MR images of 10 horses (8 gelding, 2 mare, 4-7 years old) previously published in Graham et al. (2020). Here, sagittal plane T2W images were used to calculate the articular surface normal of the lateral proximal sesamoid bone (see Fig. S2). The extension angle between the long axis of the third metacarpal bone and proximal sesamoid bone surface normal during stand was 106.9 ± 3.8° (Fig. S2). The metacarpal-sesamoid extension angle during gallop was then interpreted as 132.9° (106.9° + 26°) (Fig. 3c).

The congruent mesh blocks were assigned an elastic modulus and Poisson’s ratio of 2.5 GPa and 0.4 respectively to reflect polymethyl methacrylate. The proximal end of the metacarpal bone was tied to a single control node with translation degrees of freedom constrained (Fig. 3b). Translation normal to the proximal phalanx block was constrained, as well as axial rotation (Fig. 3b). Abaxial translation of each proximal sesamoid block was constrained to simulate tension of the intersesamoidean ligament. A displacement-controlled boundary condition was then applied to the proximal sesamoid block at an angle of 132.9° to the long axis of the third metacarpal (Fig. 3b-c). A monotonic displacement of 1.25 mm was applied at a displacement rate of 5 mm/min.

FE-simulations were conducted in Abaqus (2016a, Dassault Systèmes) where the material model in section 2.5.1 was incorporated as a user defined material subroutine (UMAT). For each metacarpal, the lateral and medial condyles were tested independently to compare relative differences in whole bone mechanical properties between each side. First, a QCT-derived BV/TV only material definition (*D*^pex^ = 0) was simulated for each limb. We then repeated the simulations with the updated MRI-derived damage surrogate (*D*^pex^ > 0) assigned to the elements that comprise the segmented palmarodistal dense bone region. The hexahedral FE-models (n=40) of the condylar compression of each metacarpal were solved using domain-level parallelisation with 32 CPUs on Heriot-Watt University’s HPC facility ‘Rocks’.

### 2.6 Data analysis

The force-displacement data was extracted from the control node tied to the proximal sesamoid congruent mesh block. From the force-displacement data we evaluated the following whole bone mechanical properties; the apparent stiffness, *K_app_*, (maximum local stiffness); yield force, *F_y_*, (force where the local stiffness is reduced to 0.9 ∗ *K_app_*) (Varga et al., 2017).

Statistical analyses were performed in python with Statsmodels (Seabold and Perktold, 2010). Wherever an intra-limb comparison between the lateral and medial side was conducted, a Wilcoxon signed-rank test was used. We used regression analysis to assess the effect of the volume of the distal metacarpal, MV, and dense bone volume fraction, DBV/MV, on whole bone mechanical properties. For both tests, we assumed a significance level of *p* = 0.05.

## 3 Results

### 3.1 PseudoCT analysis

The results of the pCT analyses are summarised in Table 2. A coronal slice of the segmented dense bone pCT is shown for a single metacarpal in Fig. 4 (a figure including all limbs is presented in Fig. S3). pCT-derived BMD in the dense bone regions were lower than QCT-derived BMD in all 10 metacarpals in both the lateral and medial condyles. There was no significant difference in the pCT-BMD medians between the lateral and medial condyles. There was no correlation between the relative volume of dense bone (DBV/MV) and the pCT- BMD medians. Exclusion of the dense bone volume and subsequent pCT generation of the lateral side of the metacarpal, using the medial side as the comparative ground truth resulted in a minimum histogram intersection match score of 0.917 in any limb (Fig. S3).

**Fig. 4.**
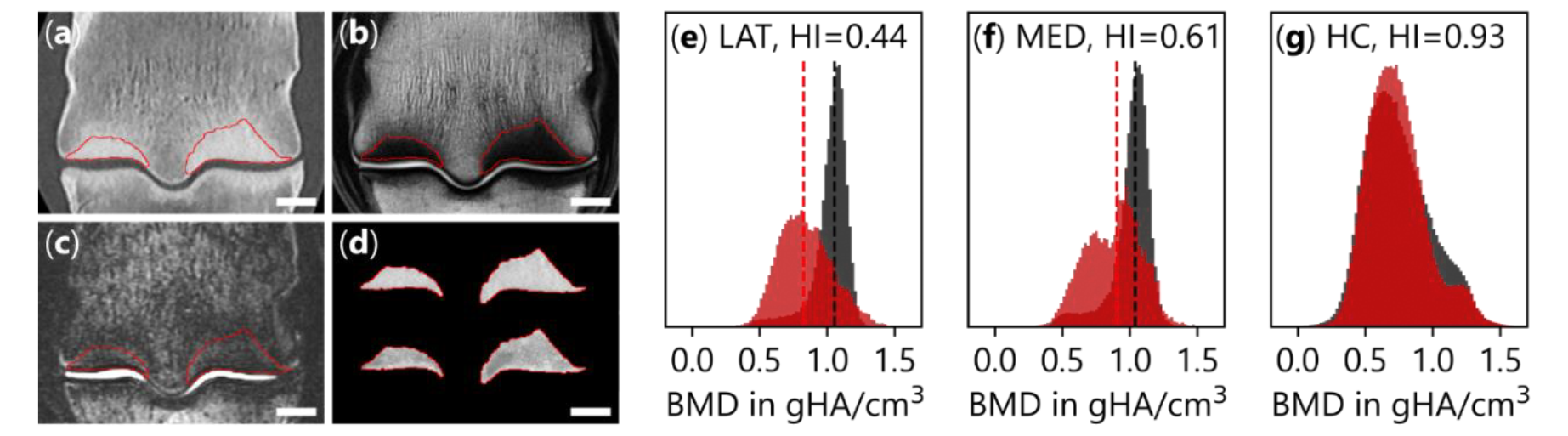
PseudoCT result of a representative limb: (a-c) Coronal slice of QCT (a), T1w (b), and STIR (c). The red line here bounds the dense bone segmentation described in section 2.4.1. Scale bar here is 10 mm. (d) The segmented dense bone where the top is the QCT, and the bottom is the pCT generated from the STIR sequence. (e-f) BMD distribution in the segmented dense bone for QCT (grey) and the pCT (red). The dashed lines signify the medians of the distributions used to quantify pre-existing damage. (g) BMD distributions of a pCT generated from the lateral side (with dense bone removed) of the metacarpal bone using the medial side as the ground truth. HI in the histogram figures denotes the histogram intersection match score.

**Table 2.** – Descriptive statistics of pCT generation results of n=10 metacarpals.

| Side | | $BMD_{QCT}$<br>in mgHA/cm<sup>3</sup> | $BMD_{pCT}$<br>in mgHA/cm<sup>3</sup> | $\rho_{QCT}$<br>in - | $\rho_{pCT}$<br>in - | $D^{pex}$<br>in - | $\kappa^{pex}$<br>in - |
| --- | --- | --- | --- | --- | --- | --- | --- |
| Lateral | Min | 1017.2 | 738.4 | 0.957 | 0.697 | 0.110 | 0.117 |
|  | Median | 1031.4 | 814.5 | 0.970 | 0.768 | 0.203 | 0.227 |
|  | Max | 1047.7 | 911.6 | 0.985 | 0.859 | 0.278 | 0.326 |
| Medial | Min | 1004.4 | 667.8 | 0.945 | 0.632 | 0.059 | 0.061 |
|  | Median | 1031.7 | 827.2 | 0.970 | 0.780 | 0.202 | 0.226 |
|  | Max | 1054.3 | 953.0 | 0.991 | 0.897 | 0.353 | 0.435 |

The influence of the MRI-derived damage surrogate, *D*^pex^, on both elastic and post-yield material behaviour is shown in Fig. 5. The result in Fig. 5 is normalised to bone volume fraction (BV/TV = 1). The minimum, median and maximum detected *D*^pex^ (Table 2) reduced all components of the stiffness tensor, 𝕊, by 5.9%, 20.9% and 35.3% respectively. Similarly, the minimum, median and maximum isotropic contraction of the quadric yield surface was 5.9%, 20.3% and 35.3%.

**Fig. 5.**
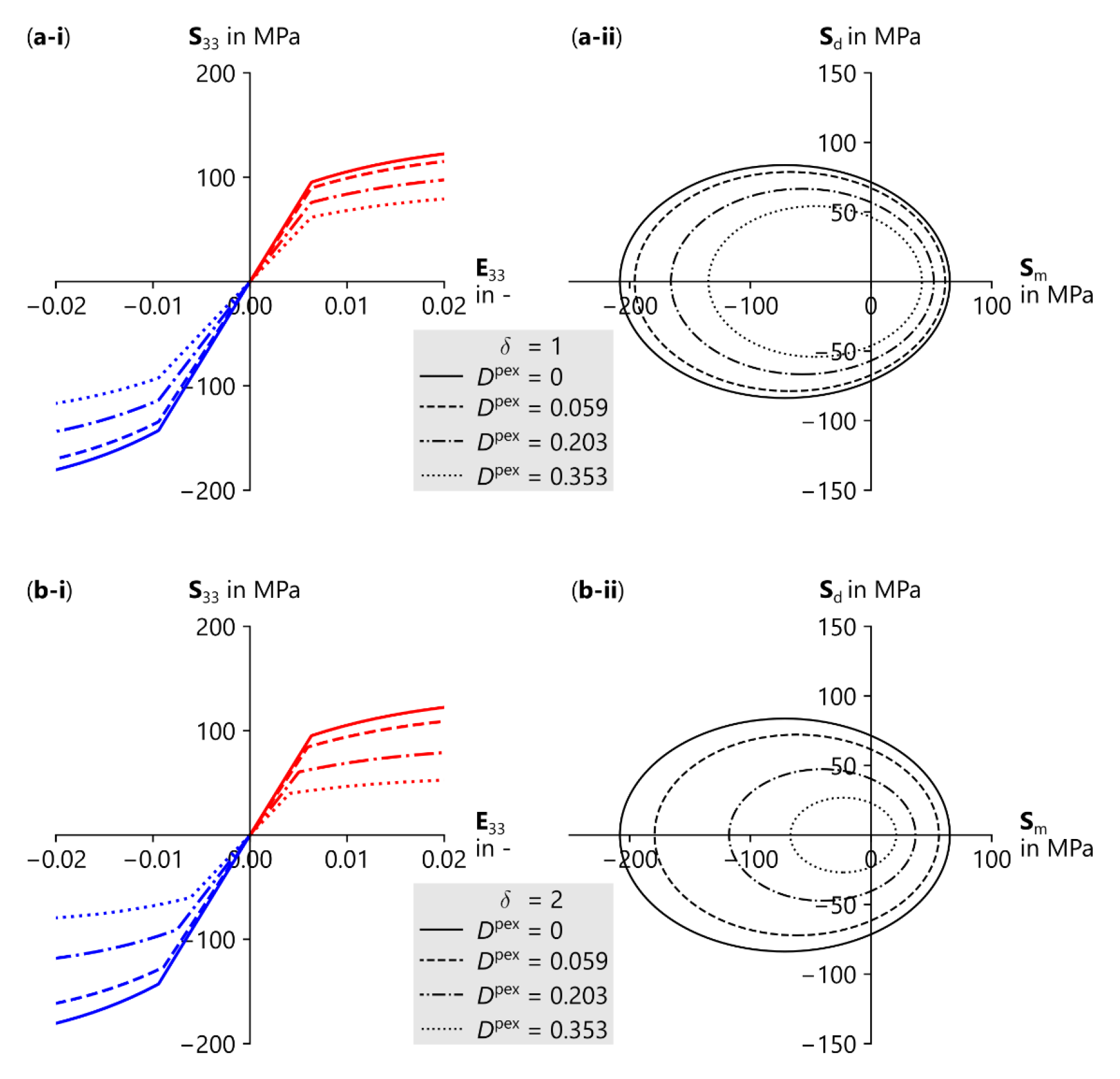
Influence of MRI-derived damage surrogate. *D*^pex^, on the material elastic and post-yield behaviour with a bone volume fraction of *ρ* = 1. (a-i) Stress-strain relationship under uniaxial compression (blue) and tension (red). (a-ii) Quadric yield surface illustrated in the mean stress vs deviator stress plane. (a and b) In all instances, the line style corresponds to the magnitude of *D*^pex^. The solid line represents the QCT-only derived behaviour, while the dashed lines illustrate the reduction in stiffness (a-i) and contraction of yield surface (a-ii) for the minimum (dashed), median (dot-dashed), and maximum (dotted) detected *D*^pex^. (b-i and b-ii) is a repeat of (a-i and a-ii), however, the interaction parameter, *δ*, in Eq. (20) is increased to *δ* = 2, which further contracts the yield surface. Note, stiffness is not influenced by the magnitude of *δ*.

### 3.2 Whole bone biomechanical properties

The QCT-derived BV/TV dependant compression of the lateral and medial condyles of each metacarpal revealed that the apparent stiffness (*K_app_*) and yield force (*F_y_*) of the lateral condyle was significantly lower than the medial condyle (*K_app_*, p<0.001; *F_y_*, p<0.001) (Fig. 6a-b). The yield force and apparent stiffness of only the lateral condyle and not the medial condyle correlated with the volume of the distal metacarpal, MV (Lateral: *F_y_*, *p* = 0.00178, R^2^ = 0.73; *K_app_*, *p* = 0.00872, R^2^ = 0.58; Medial: *F_y_*, NS; *K_app_*, NS) (Fig. 6a-b). The normalised volume of dense bone (DBV/MV) in the palmarodistal aspect of both condyles showed no correlation with either the apparent stiffness (*K_app_*) or yield force (*F_y_*) (Fig. 6e-f).

**Fig. 6.**
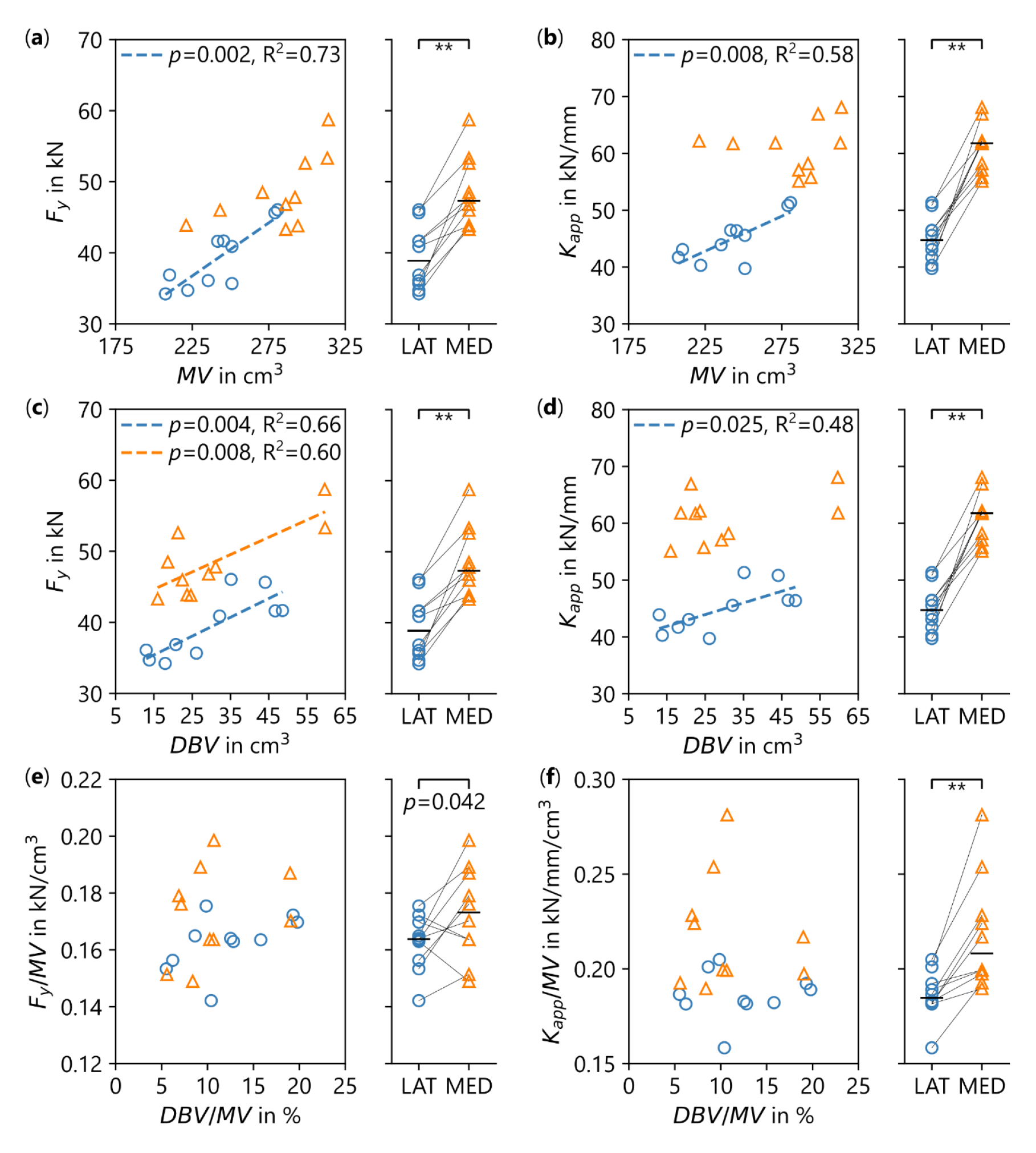
QCT-only *in silico* compression of the lateral (blue circles) and medial (orange triangles) con-dyles: (a-b) show that the yield force, *F_y_*, and apparent stiffness, *K_app_*, of the lateral condyle are dependent on the volume of the distal metacarpal, MV. For each limb, the lateral condyle was significantly (** denotes *p*<0.001) weaker and more compliant than the medial condyle. (c-d). *F_y_* and *K_app_* are corelated with the volume of dense bone DBV. (e-f) To account for the influence of metacarpal size, *F_y_*and *K_app_*were normal-ised for size by dividing by the volume of the distal metacarpal. There was no significant correlation between the normalised volume of dense bone DBV/MV and normalised yield force or apparent stiffness. When nor-malised for size, the lateral condyle remained significantly weaker (*p=*0.042) and more compliant (*p*<0.001) than the medial condyle.

The MRI-derived damage surrogate (*D*^pex^) showed no correlation with either the normal-ised volume of dense bone, DBV/MV, or the volume of the distal metacarpal, MV (Fig. 7a). Updating the material behaviour with *D*^pex^ reduced the apparent stiffness and yield force of all metacarpals for both lateral and medial condylar compression. There was no significant difference between lateral and medial relative reductions in yield force, Δ*F*_*Y*_, or apparent stiffness, Δ*K_app_* (Fig. 7b). The relative reduction in yield force Δ*F*_*Y*_ and apparent stiffness Δ*K*_*app*_ were strongly correlated with *D*^pex^ for both the lateral and medial condyle (Lateral: Δ*F*_*y*_, *p* < 0.001, R^2^ = 0.80; Δ*K*_*app*_, *p* = 0.0014, R^2^ = 0.74; Medial: Δ*F*_*y*_, *p* < 0.001, R^2^ = 0.89; Δ*K*_*app*_, *p* < 0.0011, R^2^ = 0.76) (Fig. 7b-c).

**Fig. 7.**
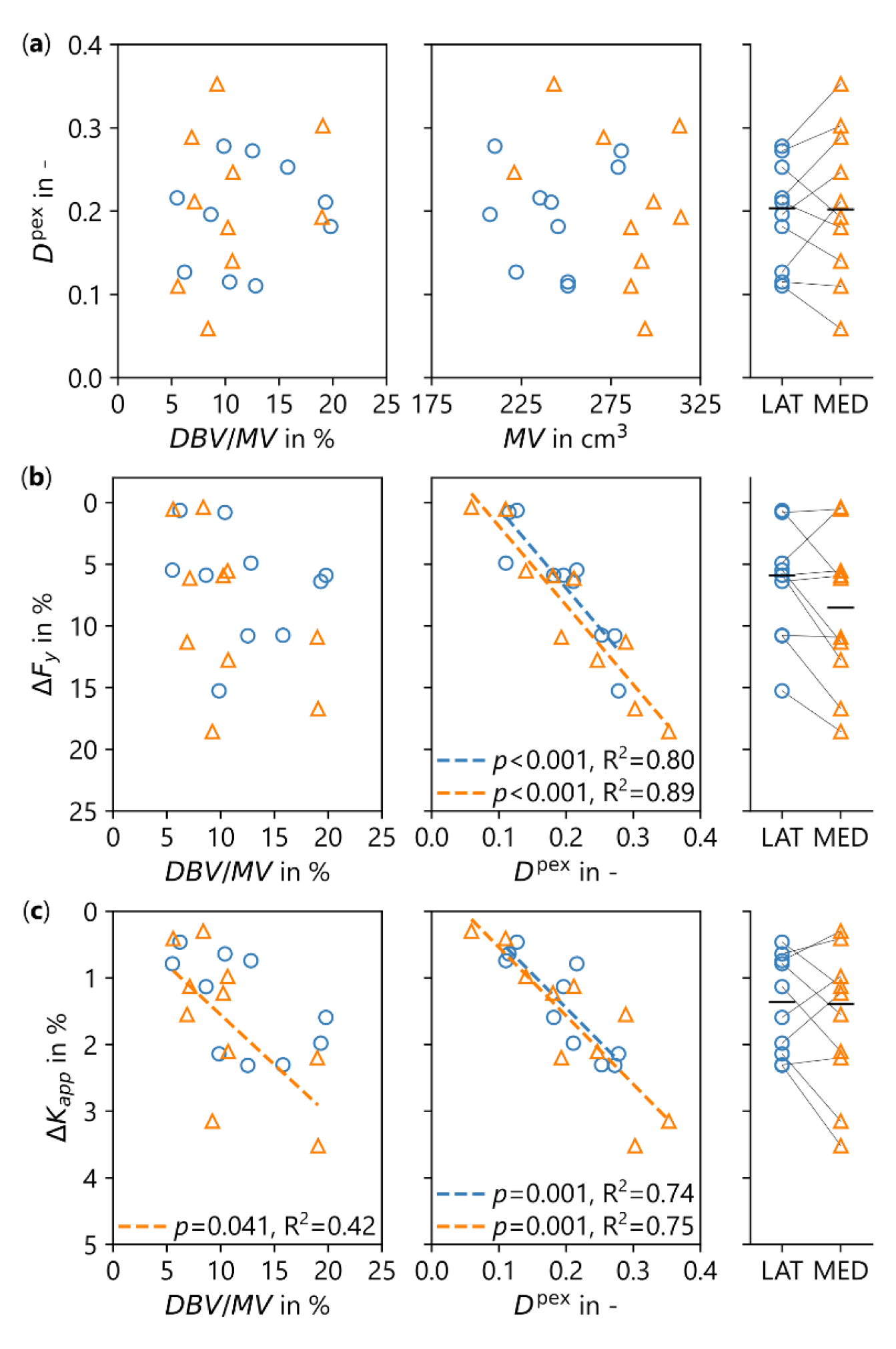
MRI-updated *in silico* compression of the lateral (blue circles) and medial (orange triangles) condyles: (a) The MRI-derived damage surrogate, *D*^pex^, shows no correlation with either the volume of the distal metacarpal, MV, or the normalised volume of dense bone, DBV/MV. The magnitude of *D*^pex^ in the lateral and medial condyle is not significant. (b) The relative reduction in yield force, Δ*F*_*Y*_, does not correlate to the normalised volume of dense bone, however, it shows good correlation with *D*^pex^. (c) Similarly, the relative reduction in stiffness, Δ*K*_*app*_, shows good correlation with *D*^pex^ and DBV/MV in the medial side of the metacarpal.

## 4 Discussion

We present in this study a methodology for updating density-based CT models with sup-plementary MRI data. We propose a method for quantifying signal increase in a fluid sensi-tive MRI sequence, which in the context of repetitive trauma, we interpret as being indicative of an underlying damage state. We then propose an update to an existing constitutive law that describes the mechanical behaviour of bone, with a pre-existing damage state variable which captures a reduction in material stiffness and strength. We then ran a comparative FE regime and show that incorporating our MRI-derived damage metric has a significant impact on bone strength which ultimately translates to an increased risk of fracture.

### 4.1.1 Multimodal imaging

Clinical QCT fails to detect microdamage *in vivo* due to the achieved volumetric resolution being several orders of magnitude larger than micron-and submicron cracks and voids. We know, however, that condylar fracture of the metacarpal is driven by a supra-physiological microdamage burden that coalesces in the dense palmarodistal subchondral bone (Muir et al., 2008, 2006; Whitton et al., 2018). The first objective of this study was therefore to identify a suitable imaging modality that is sensitive to changes in marrow composition that is provoked by repetitive loading. We chose a STIR sequence as it is sensitive to atypical water content in the adjacent bone marrow. Like QCT, this sequence does not explicitly visualise microdamage as the clinically achievable resolution is too coarse and the increased signal is none specific (Molfetta et al., 2022). However, in the context of repetitive trauma, increased signal is indicative of a stress response where edema, hemorrhage and hyperemia are present alongside microdamage (Matheny et al., 2017b; Muratovic et al., 2018; Rangger et al., 1998; Taljanovic et al., 2008). This aetiological context, coupled with evidence that exercise induced microdamage localises in the palmarodistal subchondral bone, allows us to speculate that a signal increase is representative of an underlying damage state. The stress response in bone is a transient process that reflects a spectrum of physiological changes, and we are mindful that a detected increase in signal intensity may not reflect a damaged state. However, post-mortem evaluation of racehorses that sustained a condylar fracture showed that STIR signal increase in the condyles of both a fractured limb and its contralateral nonfractured limb is significantly more common than in limbs of horses euthanised for reasons other than musculoskeletal injury (Peloso et al., 2019; Tranquille et al., 2012). This association would indicate an intrinsic link between fluid signal increases in the subchondral bone and an underlying damage state that predisposes fracture.

To quantify the STIR signal increase in palmarodistal subchondral bone, we employed a patch-based similarity search method (Andreasen et al., 2015), where the intensity of its constituent voxels is compared to the intensity elsewhere in the bone. It is important to note that this method, as used here, represents a histogram-matching approach which we used to investigate shifts in the histograms of the DBV (Fig. 4). Therefore, our results for *D*^*pex*^ represent average values on these volumes and not a location preserving voxel-by-voxel correlation that would require an establishment of a transfer function to parse MRI signal to pseudoCT without the here used registration. However, the chosen approach is sufficient to highlight the signal hyperintensity that QCT does not pick up and for which a multimodal imaging approach is beneficial.

### 4.1.2 Pre-existing damage

The BMD distribution of the MRI-derived pCT could have been directly transformed to BV/TV by the calibration law (Eq. (1)) used to transform the QCT-BMD. We chose not to incorporate the STIR signal quantification directly as a porosity, as while microdamage does ultimately manifest as a porosity, its length-scale and mechanical impact differs from the influence of structural porosity (Currey, 2002; Hernandez et al., 2014; Lambers et al., 2014; Vashishth, 2007; Wolfram et al., 2016) We instead introduce a pre-existing damage surrogate, *D*^pex^, which is a scalar state variable that captures the mechanical degradation of bone tissue incurred by the nucleation and propagation of microcracks and voids. The surrogate variable is motivated by evidence of existing and accumulating microdamage in bone (Burr et al., 1997) and a concept to include previously incurred microdamage in computational models (Mirzaali et al., 2015). *D*^pex^ is a geometric variable defined by a volume density of microcracks and other micron sized porosity and we derive it by interpreting the difference in pCT- and QCT-derived BV/TV such that the difference is interpreted as fluid occupied microscopic porosity. This direct geometric derivation infers that the signal increase in STIR sequence is directly related to microcrack density. While this may not be strictly correct, we chose this method as it allowed us to quantify the signal increase when only equipped with *in vivo* images acquired in a clinical setting. To establish an empirical relationship between the signal in the STIR sequence and the true microcrack density would require verification by imaging at micron-sized resolutions, i.e. in the size order of microcracks by staining-based histology (Arlot et al., 2008a, 2008b) or synchrotron radiation μCT (Larrue et al., 2011; Wolfram et al., 2016). Beyond the clinical scanning included in the study, we did not have access to the bones to investigate this at this stage.

We update an existing elasto-viscoplastic material model proposed by Schwiedrzik and Zysset (2013) to illustrate the impact of a potential pre-existing damage state on the mechanical behaviour of the subchondral bone. For the update, we follow Mirzaali et al. (2015) who investigated the impact of an overloading in compression on the tensile mechanical properties and vice versa. The principal idea is similar to incorporating a pre-existing damage state. As several authors have shown (Garcia et al., 2010; Mirzaali et al., 2016; Wolfram et al., 2011; Zysset and Curnier, 1996), accumulated plastic strain and bone damage are related by an exponential relationship which allows us to couple our pre-existing damage surrogate with the accumulated plastic strain additively (Eq. (14)). This additive mechanism has the effect of reducing the initial material stiffness without influencing the accumulation of damage through mechanical overloading, i.e. the pre-existing weakening and the one caused by plastic deformation are separate processes. Inclusion of our damage surrogate led to a reduction in material stiffness between 5% (14.3 GPa) and 35% (9.8 GPa). However, the maximum stiffness reduction derived here, still resulted in an apparent elastic modulus that is higher than those reported for palmarodistal subchondral bone in Thoroughbred racehorses, which ranged between 2.5 GPa to 5.6 GPa (Malekipour et al., 2018; Martig et al., 2020, 2013; Rubio-Martínez et al., 2008). Interestingly, these reported modulus values of the sub-chondral bone are 10% to 30% of that of cortical bone despite having a comparable bone volume fraction of ≈0.95. While the mineralisation of the subchondral bone is slightly lower than that of cortical bone (Boyde, 2003; Boyde and Firth, 2005), the underlying tissue level elastic modulus is only 25% lower (Doube et al., 2010). It could be inferred then that it is the structure, in which microdamage is a compounding factor, rather than the composition that is responsible for the disparity in the apparent elastic modulus. Further to reducing stiffness, microdamage also reduces bone strength (Hernandez et al., 2014; Mirzaali et al., 2015). To capture the reduction in strength, we propose an isotropic contraction of the quadric yield surface. The median contraction of 20.3% corresponds to a compressive yield stress of 113.7 MPa, which compares well to Rubio-Martínez et al. (2008), who reported a compressive yield stress of 113.3 MPa for bone specimens extracted from the palmarodistal aspect of metacarpals with subchondral bone disease. Ultimately, the material parameters *k_pex_* and *δ* (see Eq. (15) and Eq. (20)) that control *D*^pex^ and the strength reduction, respectively, are likely greater than one (Fig. 5) and therefore need to be experimentally identified.

### 4.1.3 Whole bone *in silico* monotonic strength test

Pair-wise intra-limb comparison of condylar stiffness and strength between the lateral and medial condyles revealed the medial condyle is significantly stiffer and stronger than the lateral condyle (*p* < 0.001). We speculate that this is primarily attributed to the medial condyle of all metacarpals included in our study being larger than their respective lateral condyle. This increased size, which translates to an increase in stiffness and strength, is likely a response to the higher load that is transferred to the medial condyle (den Hartog et al., 2009). When condylar stiffness and strength were normalised by the volume of the distal metacarpal, however, the lateral condyle remained weaker and more compliant than the medial condyle (Fig. 6e-f). This may explain the disproportionate incidence of lateral condylar fractures compared to medial condylar fractures in the third metacarpal bone of Thoroughbred racehorses (Parkin et al., 2004). The FE setup does not consider the morphology of the lateral and medial proximal sesamoid bones. Harrison et al. (2014) show that contact pressure is primarily localised to the parasagittal groove as opposed to being evenly distributed across the articular surface of the condyle as we model. This heterogeneous loading may influence condylar strength, especially if an underlying osteolytic lesion is present in the parasagittal groove. We chose our method as the loading was independent of joint morphology allowing for an anatomically normalised measure of condylar stiffness and strength between limbs. The localisation of accumulated damage in the parasagittal groove (Fig. 8) would indicate that the FE-setup sufficiently captures the mechanical loading of distal metacarpal, as this is where subchondral microdamage propagates to condylar fracture. Improvements could be made by developing limb-specific joint loading models similar to that proposed by Harrison et al. (2014).

**Fig. 8.**
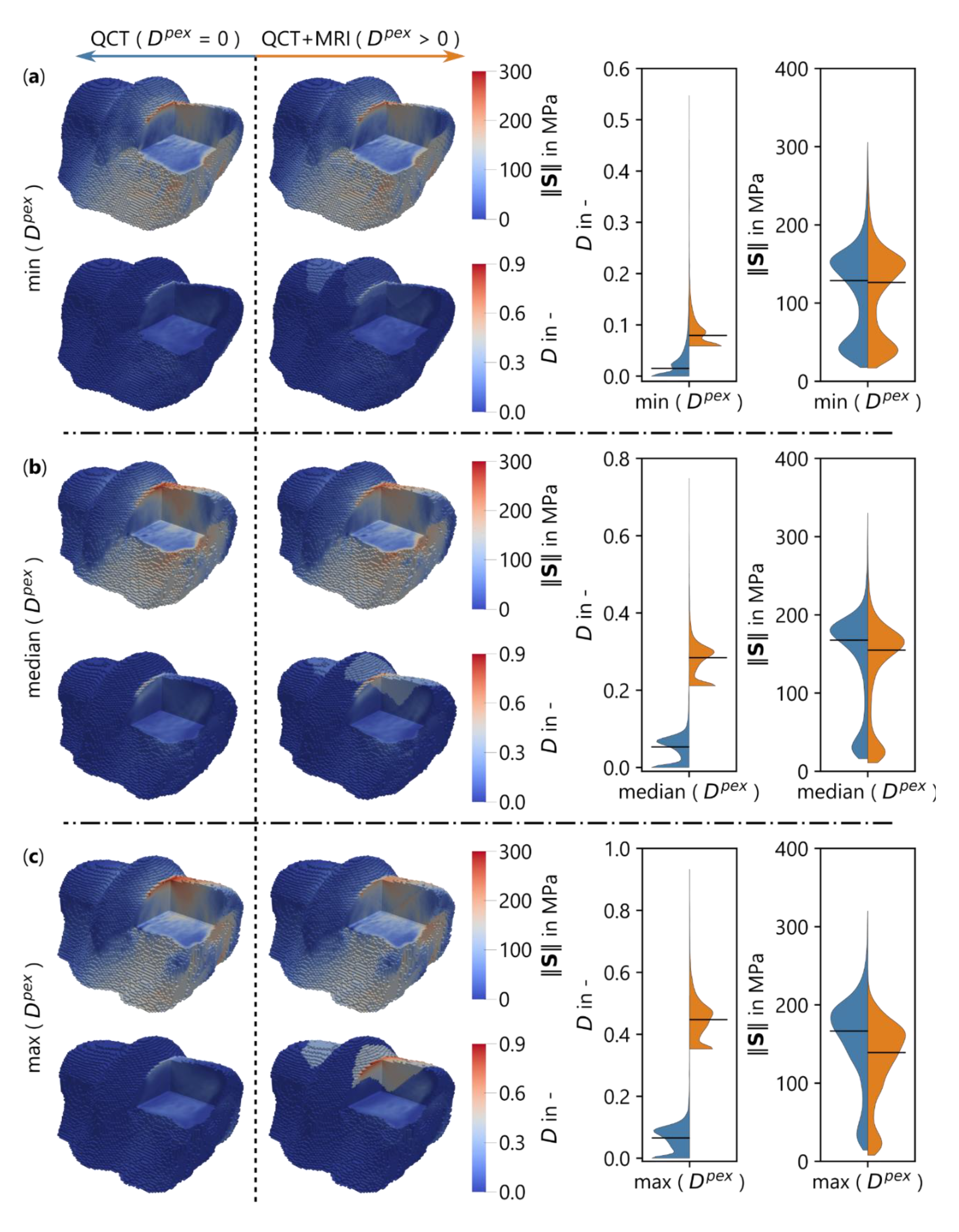
Stress norm and damage distributions in the distal metacarpals for three exemplary limbs which exhibit the minimum (a), median (b), and maximum (c) detected *D*^pex^. Within each limb case, left of the vertical dashed line is the QCT-only distributions, and right is the QCT updated with the pre-existing damage surrogate, *D*^pex^. The violin plots in each row show the distribution of damage and stress norm in the segmented dense bone region, where, left (blue) is the QCT-only and right (orange) the QCT updated with *D*^pex^. The black horizontal lines are the distribution medians.

The QCT-derived FE modelling (*D*^pex^ = 0) alone neglects the possible influence of pre-existing damage or microporosity which may lead to an overestimation of the actual mechanical competence of the distal third metacarpal bone. For example, Whitton et al. (2018) show that the density of subchondral microcracks in horses in race training is higher than in horses at rest, despite exhibiting similar subchondral bone volume fraction. In this instance, the clinical QCT would only return the subchondral bone density and overlook a weakening incurred by the underlying microdamage. We hypothesised prior to analysis that the amount of detected *D*^pex^ would correlate to the anatomically normalised measure of subchondral densification (DBVP) in the hope that this would be an accessible clinical marker of the amount of accumulated microdamage in horses in training. Our hypothesis was, however, falsified. The absence of a correlation may support the notion that subchondral densification is protective (Martig et al., 2020) and the magnitude of *D*^pex^ may then indicate case-by-case instances where this protective mechanism is overwhelmed. Our results suggest that assessing the mechanical competence, including *D*^pex^, requires multiple imaging modes and the modes we investigated appear to be good candidates. If the methodology could be corroborated by experimental validation, for example via paired multiscale mechanical and imaging techniques, we could use it to inform clinicians when to cease high-intensity exercise to limit the risk of fracture.

### 4.1.4 Study limitations

With the limited number of specimens available and the lack of experimental validation, care must be taken with regards to interpretation of the results. An influential limitation here is the lack of training history for the horses included in the study. If horses were at rest at the time of euthanasia, the root cause of any MRI signal abnormality may have resolved and either the signal abnormality is persistent and reflects no underlying tissue damage or simply may not be present. An advantage to the imaging-based approach is the ability to perform a longitudinal study following the evolution of subchondral bone changes throughout a training regimen, whereas higher resolution techniques, i.e., histology or µCT are constrained by an end point that is euthanasia. Further limitations include the lack of experimental validation of the FE modelling, which limits any direct application. However, the modelling procedure, without *D*^pex^ and *κ*^pex^, has been extensively validated in studies on bones from different anatomical locations as well as different load cases (Dall’Ara et al., 2010, 2012, 2013; Pahr et al., 2012; Schwiedrzik & Zysset, 2013; Varga et al., 2010, 2016; Wolfram et al., 2010). The scope of limb specific FE-modelling was comparative in nature, with the aim of investigating the relative impact on stiffness and strength when updated with the pre-existing damage surrogate. This comparative scope extends to the potential limitation of parameterising the material model from literature. While the parameterisation needs to be derived experimentally, it was consistent in both states before (*D*^pex^ = 0) and after (*D*^pex^ > 0) the MRI update. Irrespective of the limitations in the present study, we believe the results of this study illustrate the potential value of MR imaging in the detection of bone quality metrics which contribute to bone fragility that are independent of bone mass.

## 5 Conclusion

We here demonstrate the need for clinical imaging methods that can provide information on bone quality beyond CT-derived bone mass and architecture. We propose using MRI as a complementary modality, as it can identify bone marrow abnormalities that are indicative of underlying damage, which are not detectable through clinical CT imaging. Our method for quantifying the arbitrary signal intensity in an STIR sequence allowed us to quantify relative increases in signal intensity in the subchondral bone of the distal third metacarpal bone. Using this quantification, we derived and incorporated a damage surrogate into an updated constitutive model and demonstrated the impact of a pre-existing damage state on bone strength and stiffness that would otherwise be unattainable by clinical CT alone. As the degree of microdamage accumulation is influenced by a multitude of factors, such as age, disease, medication and exercise activity, the proposed methods may help to refine patient-specific material representations for bone tissue which may be beneficial for improved diagnostics and therapy beyond the equine model on fracture risk assessment in human diseases such as osteoarthritis, bone cancer, or osteoporosis.

## 6 Declaration of competing interest

None of the authors have a conflict of interest with respect to the work reported in the study.

## 7 CRediT authorship contribution statement

**Samuel McPhee:** Formal analysis, Investigation, Methodology, Visualization, Writing - Original Draft. **Lucy Kershaw:** Data Curation, Investigation, Writing - Review & Editing. **Carola Daniel:** Data Curation, Investigation, Writing - Review & Editing. **Marta Peña Fernández:** Methodology, Writing - Review & Editing. **Eugenio Cillán-García:** Resources, Data Curation, Funding acquisition, Writing - Review & Editing. **Sarah E Taylor:** Conceptualization, Funding acquisition, Resources, Supervision, Writing - Review & Editing **Uwe Wolfram:** Con-ceptualization, Funding acquisition, Resources, Supervision, Writing - Original Draft, Writing - Review & Editing.

## 8 Acknowledgements

This work was supported by a Leverhulme Trust Research Project Grant (RPG-2020-215) and the Engineering and Physical Sciences Research Council (EP/P005756/1) to Uwe Wolfram, as well as the Horseracing Betting Levy Board (VET/CS/027) and Siemens Project (IPA 42) to Sarah E Taylor.

## 10 Supplementary Materials

**Fig. S1.**
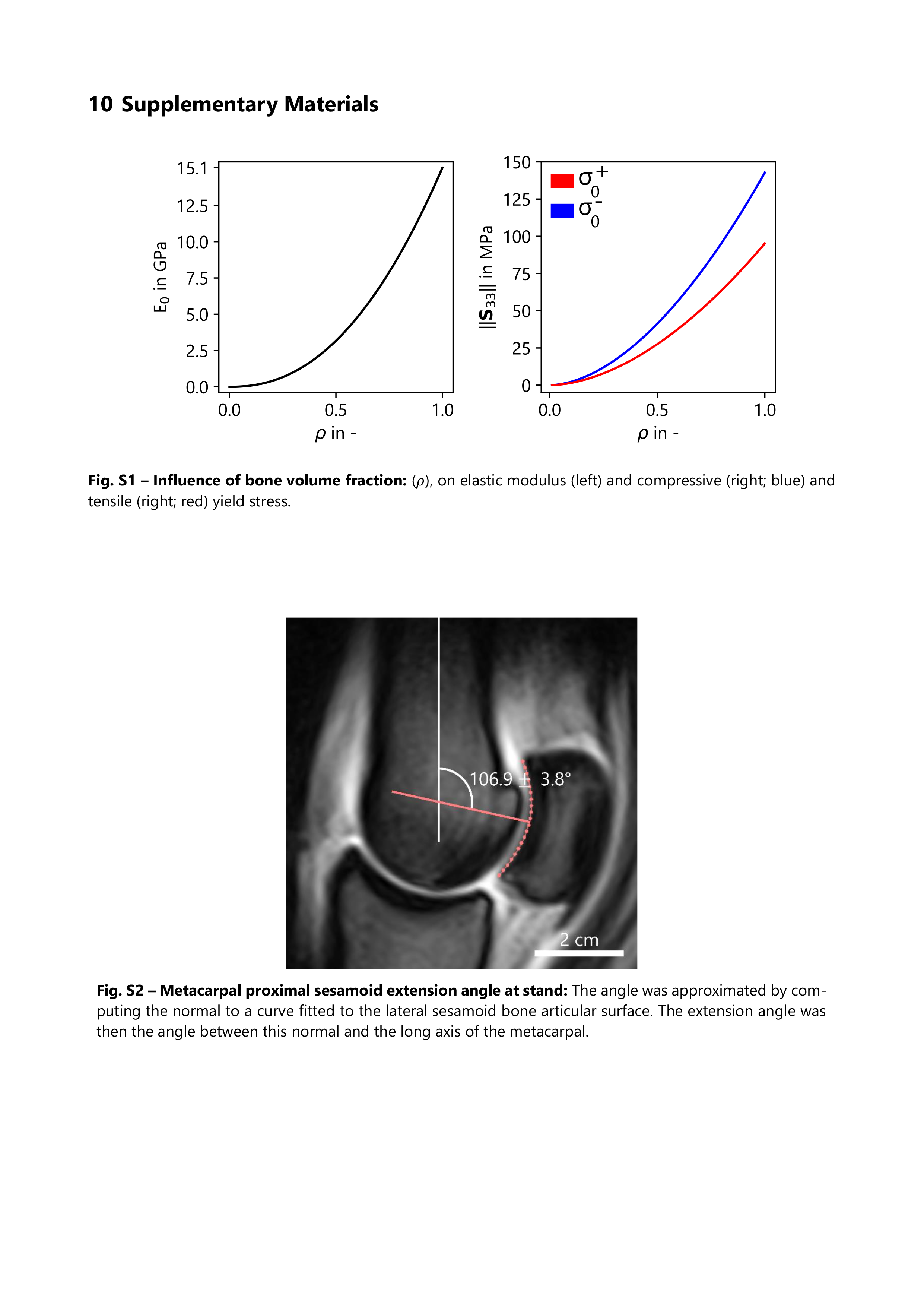
Influence of bone volume fraction: (*ρ*), on elastic modulus (left) and compressive (right; blue) and tensile (right; red) yield stress.

**Fig. S2.**
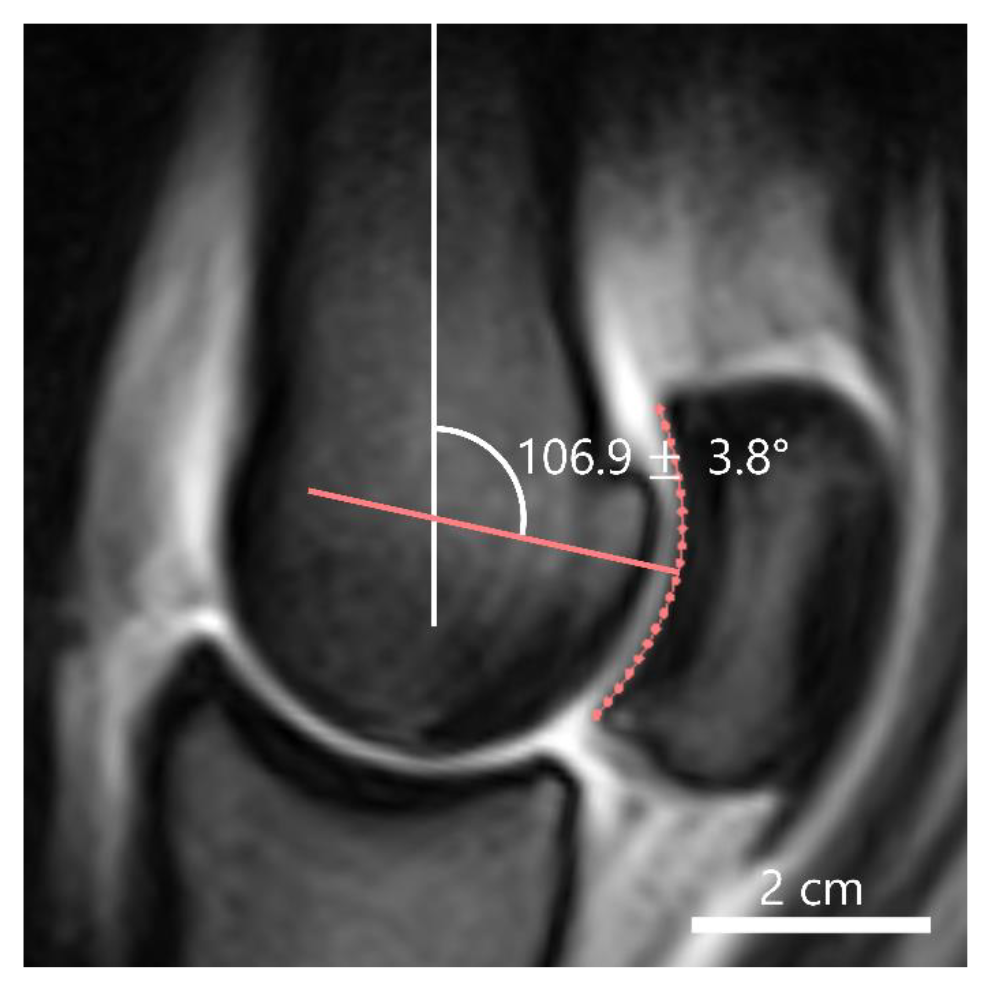
Metacarpal proximal sesamoid extension angle at stand: The angle was approximated by computing the normal to a curve fitted to the lateral sesamoid bone articular surface. The extension angle was then the angle between this normal and the long axis of the metacarpal.

**Fig. S3.**
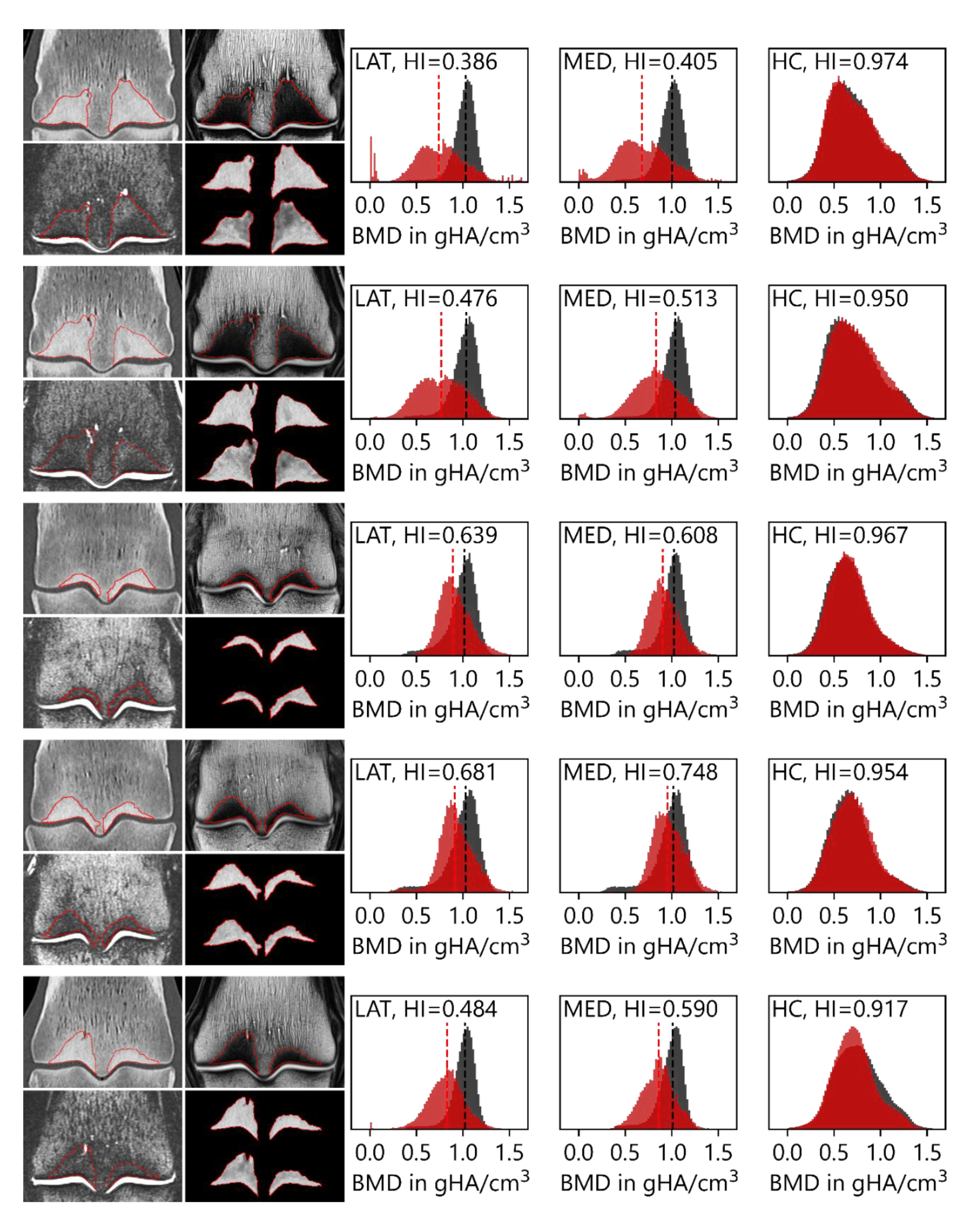
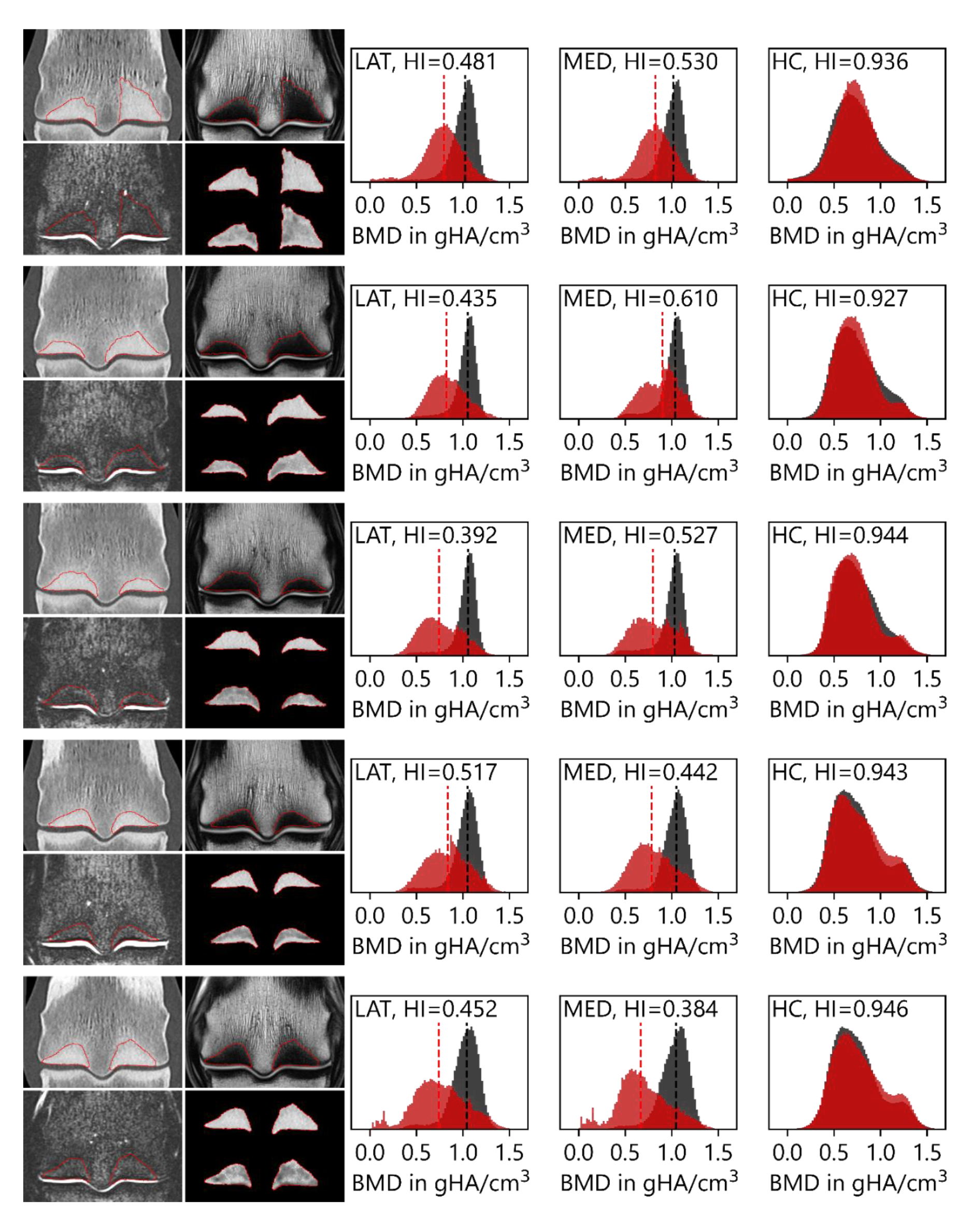
PseudoCT results for n=10 metacarpals: The image panels correspond to coronal slices of QCT (top left), T1w (top right), and STIR (bottom left). The red lines here bound the dense bone segmentation described in section 2.4.1. Scale bar here is 10 mm. (bottom left) The segmented dense bone where the top is the QCT, and the bottom is the pCT generated from the STIR sequence. The first two histograms depict the BMD distribution in the segmented dense bone for QCT (grey) and the pCT (red) in the lateral (LAT) and medial (MED) sides. The dashed lines signify the medians of the distributions used to quantify pre-existing damage. The right histogram depicts BMD distributions of a pCT generated from the lateral side (with dense bone removed) of the metacarpal bone using the medial side as the ground truth. HI in the histogram figures denotes the histogram intersection match score.

## Notes

### Competing Interest Statement

The authors have declared no competing interest.

